# CryoViT: Efficient Segmentation of Cryogenic Electron Tomograms with Vision Foundation Models

**DOI:** 10.1101/2024.06.26.600701

**Authors:** Sanket R. Gupte, Cathy Hou, Gong-Her Wu, Jesús G. Galaz-Montoya, Wah Chiu, Serena Yeung-Levy

## Abstract

Cryogenic electron tomography (cryoET) directly visualizes subcellular structures in 3D at the nanometer scale. Quantitative analyses of cryoET data can reveal structural biomarkers of diseases, provide novel mechanistic insights, and inform the effects of treatments on phenotype. However, existing automated annotation approaches primarily focus on localizing molecular features with few methods accurately quantifying complex structures such as organelles. We address this challenge with CryoViT, a paradigm shift from traditional convolutional neural networks that leverages vision transformers to enhance the segmentation of large pleomorphic structures that can occupy almost the entire field of view in high-magnification images, such as mitochondria. CryoViT is powered by a large-scale vision foundation model and overcomes limitations of popular U-Net based methods, particularly when training data are scarce. We demonstrate the efficacy of CryoViT on a large cryoET dataset of neurons differentiated from iPSCs derived from Huntington disease (HD) patients and cultured HD mouse model neurons.

---

Cryogenic electron tomography (cryoET) is an advanced imaging technique that creates 3D density maps of subcellular components, offering detailed insights into the structures of macromolecules, organelles, and cytoskeletal elements in their native context across different length scales ^1^. Extensive image processing of subvolumes through subtomogram averaging (STA) ^2,3^ allows for the visualization of macromolecular complexes such as ribosomes bound with drugs *in situ* at near-atomic resolution ^4^. Improvements in cryogenic preservation of biological samples offer the potential to study a wide range of complex biological systems across multiple cell types and organisms, providing the opportunity to reveal ultrastructural changes in different physiological states. Such direct 3D visualization of subcellular features can transform our understanding of the mechanisms underlying diseases ^5^. Indeed, cryoET is becoming increasingly relevant in biomedical applications ^6^, including the characterization of therapeutic agents such as nanoparticles ^7^. As a concrete example, we previously used cryoET to quantify the volumes and distributions of mitochondrial granules in neurites of human Huntington’s disease (HD) patient iPSC-derived and mouse model neurons with varying trinucleotide repeat lengths, thereby identifying novel cellular biomarkers of HD ^8^. Additionally, such measurements corroborated that knockdown of the PIAS1 gene ameliorates HD phenotypes in neuronal mitochondria ^9,10^, suggesting that cryoET may be useful to evaluate the effects of potential therapeutic interventions. Similar applications of cryoET have explored the ultrastructure of primary cilia ^11^, cells from patients with Leigh Syndrome ^12^, platelets from patients with acute myeloid leukemia ^13^ and ovarian cancer ^14^, as well as other human and mouse cancer models ^15^. In spite of strategies to speed up cryoET data processing and analysis ^16,17,18,19,20,21,22,23^ , manual annotation of complex, pleomorphic features in tomograms at scale is untenable ^24^, even if using interpolation strategies to increase efficiency ^25^. Therefore, building automated systems capable of mining complex feature patterns across hundreds of tomograms remains a fundamental challenge to meet the increasing demand in this rapidly evolving field.

To develop accurate and efficient models for quantifying subcellular structures across heterogeneous and challenging conditions (*i.e.*, cryoET tomograms of thick specimens, resulting in images with high noise, low contrast, contrast transfer function artifacts, missing wedge and tomographic reconstruction artifacts, etc.), we leverage vision foundation models ^26–28^, which are large-scale, deep neural networks pre-trained on hundreds of millions of natural images. They have demonstrated excellent performance on downstream vision tasks, even with limited training data, and are based on the Vision Transformer ^29^ (ViT) architecture, which captures the global context of features in an image as opposed to only focusing on local patches modeled by convolution layers. These properties make foundation models an excellent fit for accurately segmenting large subcellular features in cryoET data. We introduce CryoViT, a novel approach powered by a vision foundation model with over a billion parameters, which accurately segments pleomorphic 3D structures across diverse sample conditions. Our model is trained in a semi-supervised fashion and can learn from sparsely labeled slices, significantly reducing the manual effort involved in training the model and in cleaning up false positives post-annotation. CryoViT is a method to not just leverage foundation models but address the larger problem of building label-efficient models for segmentation of organelles in a near-native state, which can generalize robustly across variations in sample conditions, thereby enabling quantitative analyses at scale.

We demonstrate our approach with a curated dataset of tomograms that have been labeled to identify mitochondria. The tomograms correspond to neurites in neurons differentiated from induced pluripotent stem cells (iPSCs) derived from HD patients, and in cultured primary HD mouse model neurons ^8^. Our dataset spans 256 tomograms representing 11 different sample conditions that exhibit varied mitochondrial phenotypes. Mitochondria are not only a challenging target for deep learning models due to their pleomorphism and large size relative to the field of view in high-magnification cryoET experiments, but they also play a vital role in cellular processes and are implicated in a variety of pathological conditions ^30,31^, including neurodegeneration ^32–34^. This makes them a target of great interest to the broader biomedical scientific community. We use this dataset to benchmark both the segmentation quality and label efficiency of CryoViT and demonstrate that it outperforms the leading 3D U-Net based methods ^35,36^. Our open source software package can be used to segment mitochondria or build custom models for other structures, accelerating the discovery of novel insights by enabling quantitative analyses and rapid mining of features across a wide spectrum of cryoET datasets.

## RESULTS

### Robust feature extraction with Foundation Models

CryoViT is a deep learning method that combines a vision foundation model (DINOv2 ViT-g/14) for feature extraction from tomograms, and a 3D convolutional neural network (CNN) that upscales these features to segmentation masks (**Figure 1**). While foundation models like DINOv2 ^28^ excel across various vision tasks with minimal labeled data, their evaluation primarily relies on natural images, leading to potential challenges when applied to specialized domains such as unstained biological samples and low contrast cryoET images due to limited representation in training data.

**Figure 1.**
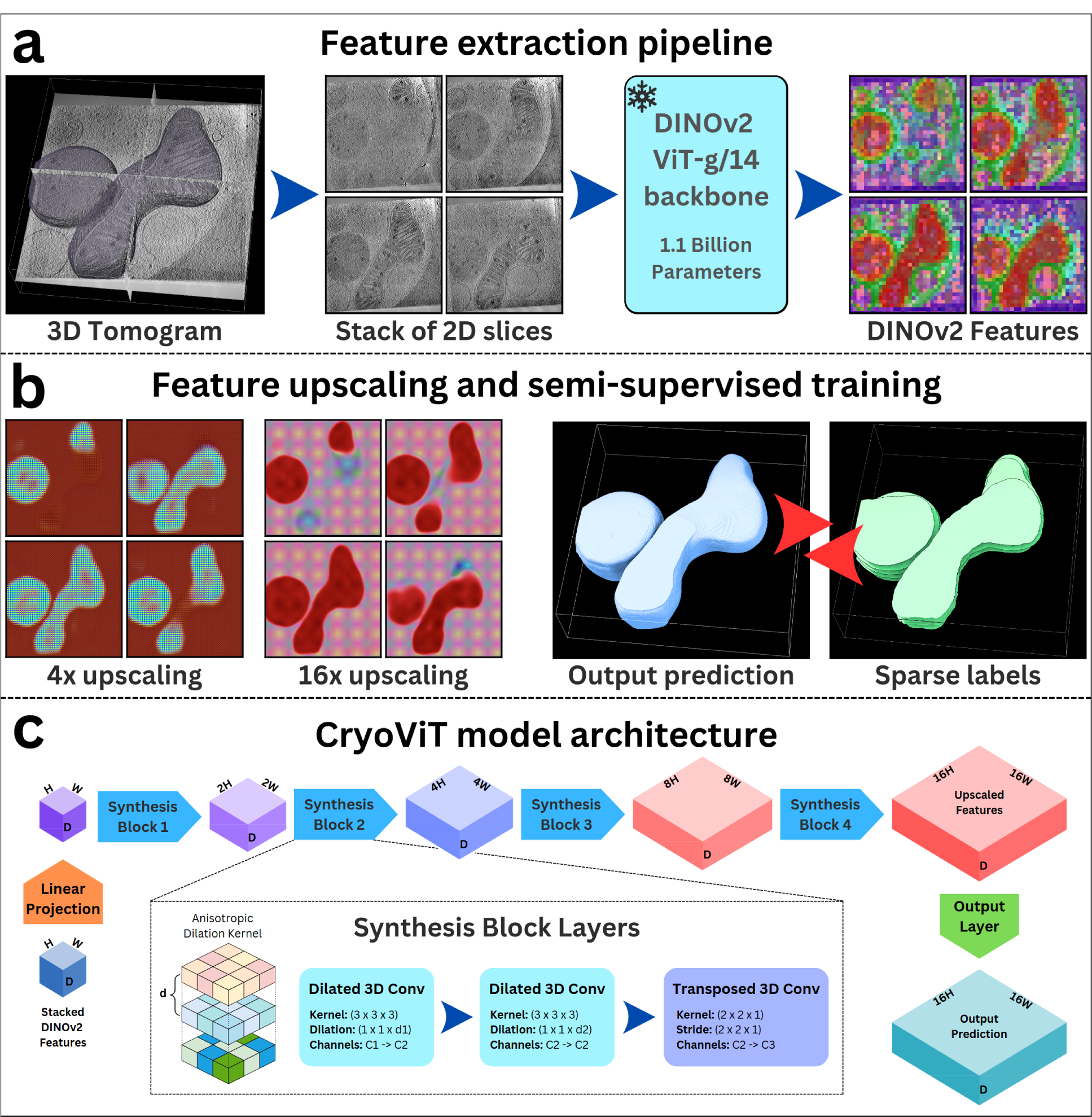
CryoViT model architecture and workflows. **a.** Feature extraction with a pre-trained vision foundation model: the 3D tomogram is split into 2D slices along the depth dimension (z). These grayscale images are fed into a DINOv2 ViT-g/14 foundation model to generate patch features (visualized using UMAP), which are stacked together to create a 3D feature grid. **b.** CryoViT’s semi-supervised training pipeline: the resulting feature grid is progressively upscaled by the CryoViT model to generate a segmentation prediction for each voxel. The model is trained on a sparse subset of labeled slices in a semi-supervised fashion. **c.** CryoViT model architecture: a pipeline of four synthesis blocks upscales the feature grid by a factor of 2 at each stage. Each block consists of a group normalization layer (omitted for clarity), two anisotropic dilated convolution layers, and transposed convolution layer to upscale in the height (y) and width (x) dimension. The 3 cartesian dimensions (x, y, z), are respectively labeled in the blocks as “W” (width), “H” (height), and “D” (depth).

Our preliminary experiments (**Figure 2**) demonstrate DINOv2’s capability to generate rich representations of cellular cryoET data without fine-tuning, underscoring its potential as a robust feature extractor for subcellular structures, including mitochondria. Unlike traditional CNNs, ViTs divide input images into non-overlapping patches called tokens and model relationships between all token pairs. This enables ViTs to capture global context information from the entire image, making them suitable for modeling structures across tomograms and their interactions, unlike CNNs limited by local receptive fields. By leveraging the full context of input images, ViTs enhance the modeling of complex structures in cryoET datasets.

**Figure 2.**
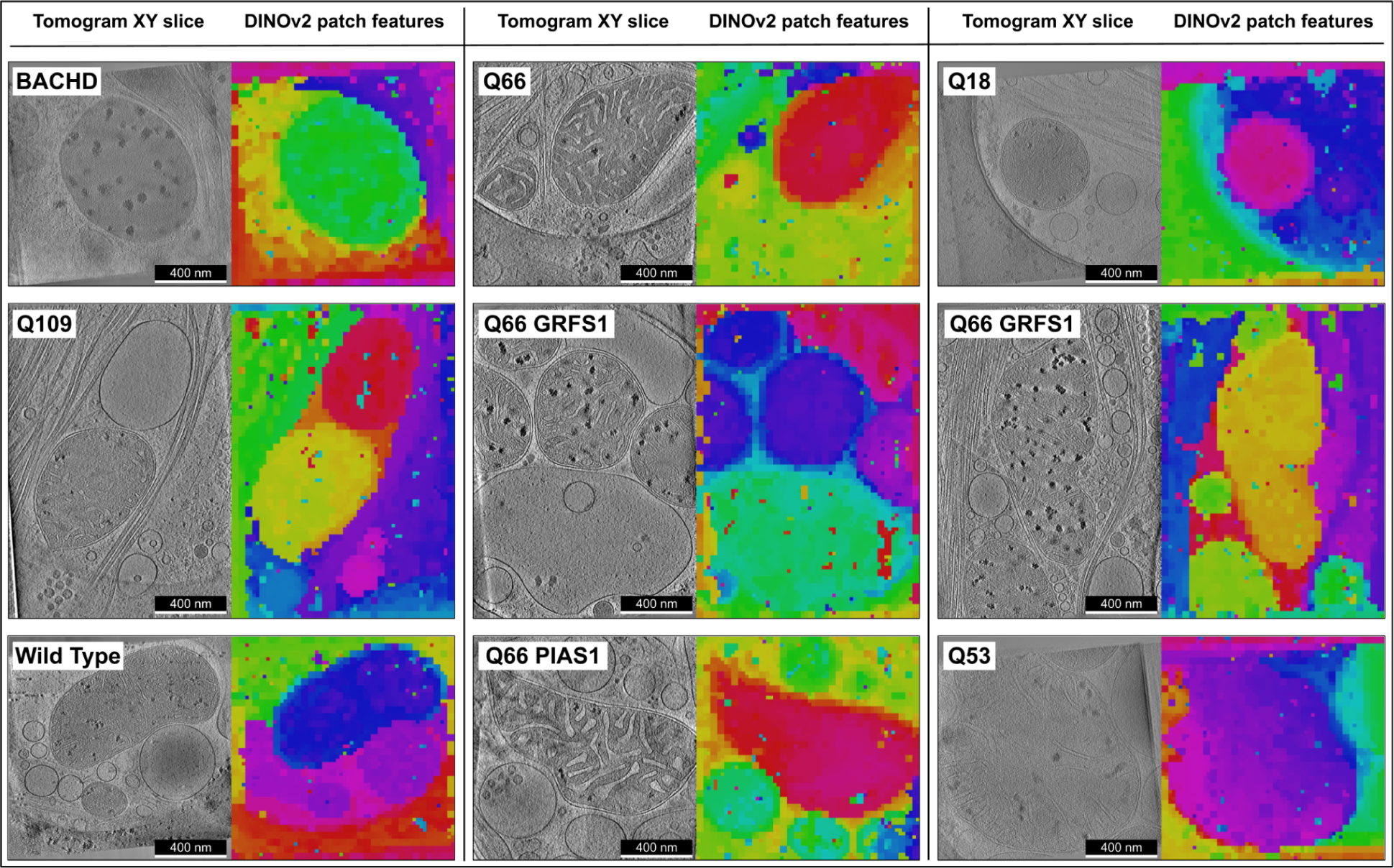
DINOv2 ViT-g/14 foundation model trained on natural images is surprisingly effective at extracting patch features from cryoET tomographic slices. Representative xy slices (grayscale images on the left side of each panel) from selected examples of reconstructed cryoET tomograms of neurites in HD patient iPSC-derived (Q18, Q53, Q66, Q109, Q66 GRSF1, Q66 PIAS1) and mouse model (wild type, BACHD) neurons, exhibiting mitochondria and other subcellular structures. The corresponding representations generated by the DINOv2 vision foundation model (color images) are shown on the right side of each panel. Patch features are extracted from each slice of a tomogram and colored using UMAP. These features display a remarkable degree of coherence with an implicit grouping of mitochondria and other subcellular structures such as vesicles. The groupings are represented by distinct color hues (arbitrarily assigned) corresponding to similar patches in latent space, highlighting the rich information stored in these types of representations. Each color in the patch feature maps corresponds roughly to the same object within each image. This process is carried out independently for each tomogram, resulting in varying hues across the different panels. Scale bars = 400 nm.

### CryoViT: modeling CryoET data with Vision Transformers

There are multiple inherent difficulties in building generalizable systems for cryoET that can accurately segment objects with limited manual annotation. The missing wedge in tilt series leads to anisotropic tomograms, and the resulting reconstruction artifacts cause a loss in resolution along the Z-axis (perpendicular to the detector). Low signal-to-noise ratio and contrast further complicate feature identification. Overcoming these challenges requires going beyond slice-based 2D annotation approaches and integrating information across the Z-axis, especially when considering organelles that can span a large fraction of the tomogram, such as mitochondria visualized at relatively high magnification.

While ViTs offer potential solutions, adapting them for cryoET analysis is not straightforward. They only produce a grid of 2D patch-level features instead of 3D pixel-level features that can be easily decoded into segmentation masks. To address this, CryoViT first generates a 3D feature grid from Z-axis slices treated as grayscale images by passing each image through the DINOv2 ViT and stacking the resulting features together. Next, a 3D CNN with anisotropic convolution kernels ^37^ is used to simultaneously upsample patch features and aggregate information across the Z-axis. In our method, we freeze the ViT backbone and train only the 3D CNN, making it possible to leverage pre-trained foundation models with over a billion parameters while using relatively modest computational resources such as a single scientific computing workstation equipped with a GPU. The 3D CNN’s lightweight design and relatively small number of parameters accelerate training and minimize the risk of overfitting (for a detailed explanation of CryoViT’s architecture, see the Online Methods).

### Semi-supervised learning for label-efficient training

To reduce annotation costs, CryoViT is trained in a semi-supervised fashion unlike competing methods which are fully supervised ^36^. Tomograms in our dataset are ∼128 slices thick in the z-dimension, and either 512 x 512 pixels or 720 x 512 pixels in the x and y dimensions depending on the sample, making it challenging to manually label each voxel. CryoViT only requires a few (∼4%) manually labeled slices per tomogram for training (n=5 for our datasets), learning from both the unlabeled and labeled regions of the tomogram in a semi-supervised fashion using a masked loss function. We manually select five evenly spaced 2D slices for annotation per tomogram, visually ensuring that each slice contains densities attributable to mitochondria while maximizing the spacing between slices. We exploit the anisotropic resolution by identifying slices at both extremes of the Z-axis beyond which the resolution is too poor to discern any mitochondria. Slices beyond these limits are automatically labeled as background (negative references), effectively increasing our labeled data for training without additional annotation costs. This approach enhances the precision of the model by discouraging spurious predictions in these regions.

### CryoViT accurately segments mitochondria across diverse samples with generalizable performance

#### Comparison of generalization performance across samples

To evaluate CryoViT’s performance in data and label constrained domains, we benchmark it on 11 different samples of mitochondria spanning as few as 10 tomograms per sample to as many as 63 tomograms per sample as we reported previously ^8^ (**Supplementary Table 1** summarizes the data). As the standard architecture for such applications ^36^, a lightly modified 3D U-Net ^35^ serves as a baseline for comparison. The CryoViT and 3D U-Net models were both trained on identical sets of data by optimizing the Dice loss ^38^ as measured on the sparse, 5-slice ground truth annotations and negative-reference slices, and performance was measured using cross-validation (see Online Methods).

Our results (**Figure 3**) indicate that, qualitatively, CryoViT produces cleaner segmentations (**Figure 3a**), and quantitatively outperforms the 3D U-Net across samples by up to a factor of 2x or more in median Dice score (**Figure 3b**, **Supplementary Table 2**). In samples with the lowest number of tomograms (n=10 for Q53_KD, and n=10 for Q66), we observe differences in median Dice score as high 0.47 and 0.31, but this gap tends to narrow as more training data are introduced. For samples with ∼30 tomograms (n=31 for Wild Type, and n=33 for Q109), the performance gap between CryoViT and 3D U-Net remains significant, with a difference in median scores around 0.1, but this reduces to 0.03 when the number of tomograms are doubled (n=63 for Q66). Although more data might narrow this gap, collecting increasingly large datasets is challenging and may be infeasible for rare conditions, structures that are difficult to find within cells and tissues ^39,40^, or complex samples that are difficult to harvest and process.

**Figure 3.**
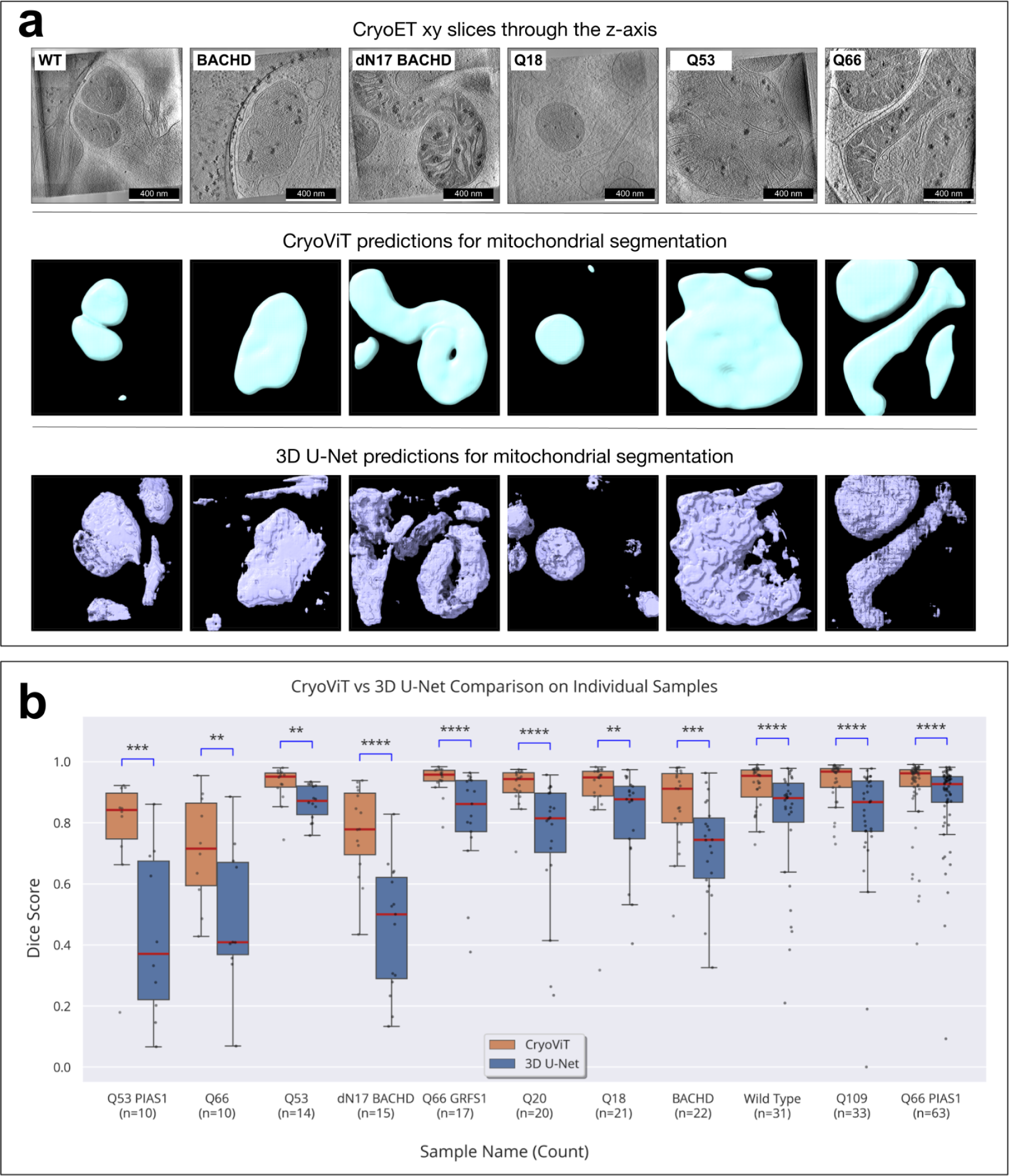
Evaluating model generalization on individual samples. **a.** Qualitative results for experiments evaluating model generalization on individual samples: comparison between CryoViT (teal) and 3D U-Net (purple) predictions on tomograms of BACHD, dN17 BACHD, Wild Type, Q53, Q66, and Q18 samples. CryoViT segmentations are smooth and continuous, closely matching the mitochondria in the corresponding tomograms (top row), even when the mitochondria span the entire tomogram. In contrast, the 3D U-Net segmentations are less accurate. Scale bars = 400 nm **b.** Quantitative results for experiments evaluating model generalization on individual samples: CryoViT (orange) outperforms 3D U-Net (blue) on all samples with higher median Dice scores and lower interquartile ranges. The red line indicates the median Dice score and scattered dots represent the Dice scores of each individual tomogram. The annotations above boxes represent the p-values for a one-sided paired significance test. ns = not significant, ****p < 0.0001, ***p < 0.001, **p < 0.01, *p < 0.05.

Qualitative observations of the predictions generated by CryoViT revealed that a Dice score greater than 0.9 represents a near perfect segmentation with only minor errors. In contrast, the predictions from the 3D U-Net exhibited gross inaccuracies, potentially missing important insights or leading to inaccurate interpretations. High-quality segmentations, such as those generated by CryoViT, can facilitate accurate downstream quantitative analyses of morphology and structure, which may be essential for feature classification and data interpretation, in turn enabling early disease diagnostics and high-resolution monitoring of disease progression ^6^.

#### CryoViT generalizes across sample domain shifts

A desirable quality for a predictive model is its robustness to domain shifts; *i.e.*, scenarios in which the distribution of data used to train the model differs from the distribution of data on which it is used to make predictions, such as when examining different samples under varied specimen preparation and data collection conditions. In this context, we explore the impact of domain shifts caused by the presence of HD phenotypes with varying severity in human and mouse model samples. Two sets of data were created: a healthy set consisting of tomograms from the Q18, Q20, and Wild Type samples, and a disease set composed of tomograms taken from HD samples without any therapeutic treatment. The mitochondria in neurites of disease samples are characterized by the presence of enlarged granules as well as altered cristae, both of which contribute to distinctly different phenotypes compared to healthy samples. This is an important case study for domain shifts because it is often easier to gather data from healthy specimens for training a model, but this model may be utilized to make predictions on diseased specimens over the course of its deployment.

We tested CryoViT’s robustness by studying domain shifts in both directions, *i.e.*, training on healthy samples and evaluating on diseased samples, and vice versa. As before, a 3D U-Net is used as a comparative baseline to contextualize the performance of CryoViT, and the models are evaluated on their ability to segment mitochondria using the Dice score. Our results (**Supplementary Table 3**) indicate a clear difference in performance between the models: CryoViT has a higher mean Dice score across all samples, highlighting its ability to generalize robustly across both domain shifts (**Figure 4a**). Interestingly, the 3D U-Net has a marginally higher median Dice score on the Q109 sample, but a handful of poor predictions lower the mean score. Though the U-Net occasionally outperforms CryoViT, such instances are the exception rather than the norm.

**Figure 4.**
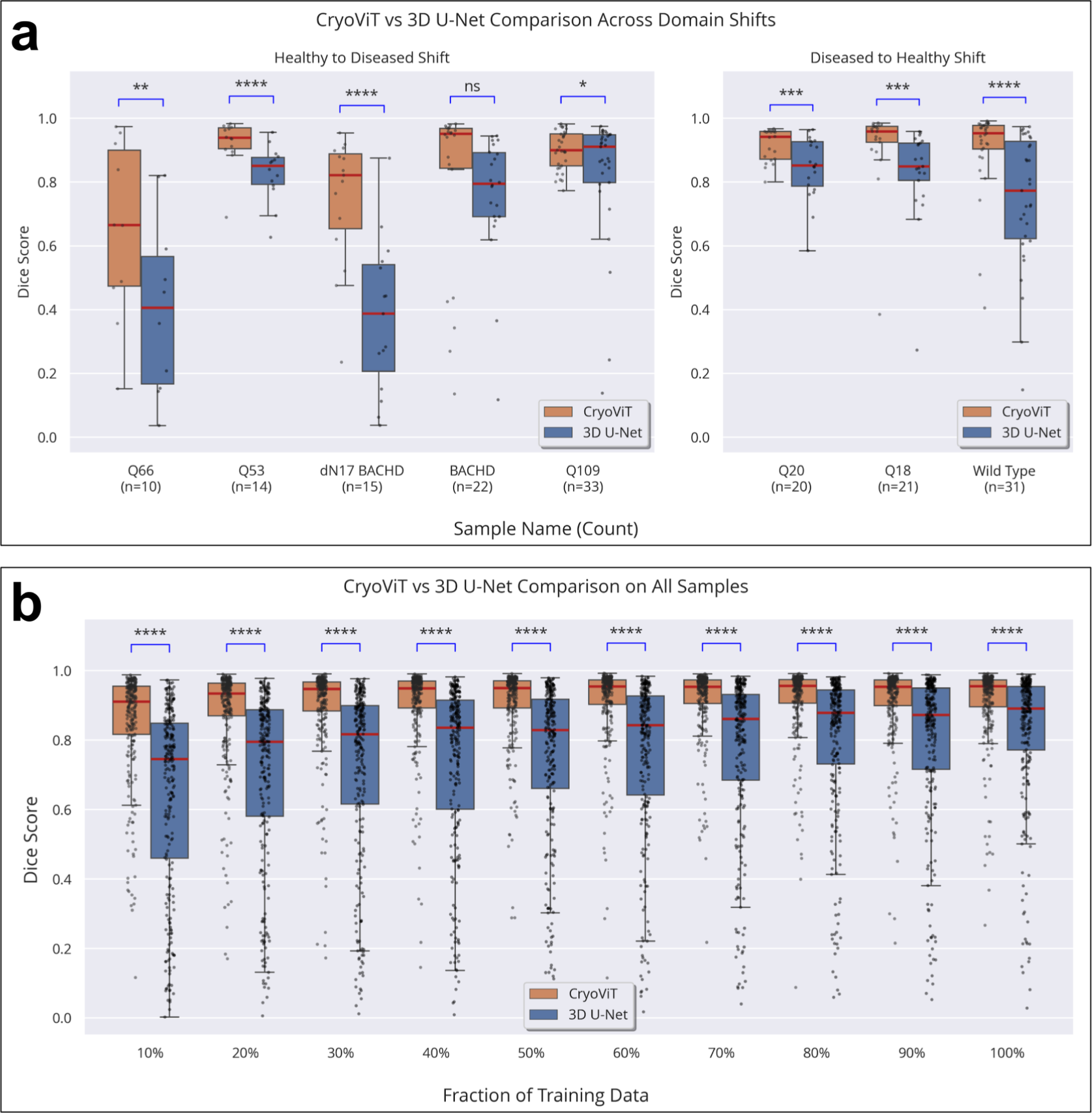
Model generalization across sample domain shifts and on unseen samples. **a.** Quantitative results for experiments evaluating model generalization across domain shifts: CryoViT (orange) outperforms 3D U-Net (blue) on nearly all samples with higher median Dice scores. The panel on the left depicts the performance of models trained on healthy samples and evaluated on diseased samples. The panel on the right depicts the performance of models trained on diseased samples and evaluated on healthy samples. **b.** Quantitative results for experiments evaluating data efficiency and model generalization on unseen samples: CryoViT (orange) outperforms 3D U-Net (blue) across all training data fractions with higher median Dice scores and lower interquartile ranges. The red line indicates the median Dice score and scattered dots represent the Dice scores of each individual tomogram. The annotations above boxes represent the p-values for a one-sided paired significance test. ns = not significant, ****p < 0.0001, ***p < 0.001, **p < 0.01, *p < 0.05.

#### CryoViT requires fewer annotations to match U-Net performance

We explored how a model’s ability to segment mitochondria from previously unseen data distributions is influenced by the number of tomograms used for training. In this evaluation setting, we employed leave-one-out cross validation at the sample level. Of the 11 different samples in our dataset, we excluded one and trained a model on the remaining 11 samples. We then calculated the Dice score for the excluded sample’s tomograms. By repeating this process for each sample, we computed Dice scores for all 256 tomograms. To study the efficiency of our models, we repeat this process by using varying fractions of the training samples (10% - 100% in increments of 10%) to train the model.

Our results (**Supplementary Table 4**) show that CryoViT models can effectively be applied to unseen samples with a modest amount of training data, requiring just 10% (27 tomograms) to achieve an impressive 0.91 median Dice score (**Figure 4b**). This performance is close to the peak Dice score of 0.96 obtained by training on 100% of the training data. In contrast, the 3D U-Net achieved a Dice score of just 0.73 with the same 10% of the training data. While this increases to 0.9 when trained on 100% of the tomograms, the U-Net fails to bridge the performance gap. Notably, this matches the performance of CryoViT trained on a tenth of the data, highlighting our method’s efficiency.

#### Dense labels minimally improve CryoViT performance

The process of manually annotating each voxel in a tomogram is extremely time consuming ^24,25^, taking anywhere between several hours to multiple days depending on the morphological complexity of the object being studied. It is impractical to scale up this laborious process to current CryoET workflows that produce hundreds of tomograms per day ^41^. Given these annotation constraints, we evaluated whether it is more effective to densely label fewer tomograms or sparsely label more tomograms. To perform this investigation, we first converted the sparse labels to dense labels using an automated procedure (see Online Methods). Next, we measured the maximum performance that could potentially be obtained by training CryoViT models on densely labeled tomograms and compared them with models trained on the same tomograms but with sparse labels (**Figure 5a**). We repeated the previous experiments evaluating mitochondria segmentation performance on individual samples as well as experiments investigating held-out performance as a function of training data fraction.

**Figure 5.**
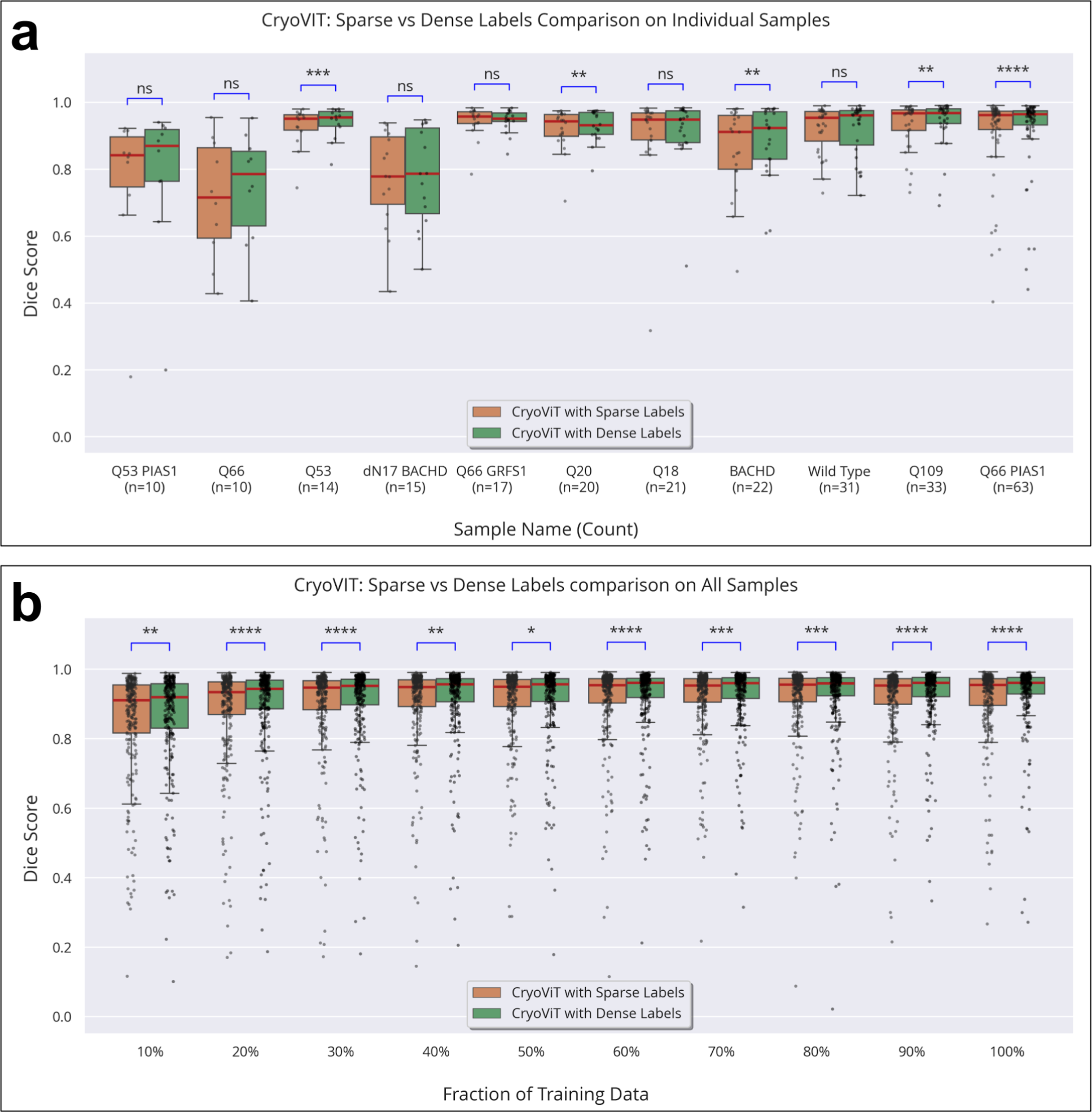
The impact of training CryoViT on densely labeled tomograms. **a.** Quantitative results for experiments comparing CryoViT variants on individual samples: In most cases, there is no significant difference between the median Dice scores of the CryoViT models trained on sparse labels (orange) vs those trained on dense labels (green). When the difference is significant, it is no more than 0.01. **b.** Quantitative results for experiments comparing CryoViT variants on data efficiency and model generalization on unseen samples: CryoViT trained on dense labels (green) outperforms the variant trained on sparse labels (orange) across all training data fractions, but the difference in mean Dice scores is just 1%-2%. The red line indicates the median Dice score and scattered dots represent the Dice scores of each individual tomogram. The annotations above boxes represent the p-values for a one-sided paired significance test. ns = not significant, ****p < 0.0001, ***p < 0.001, **p < 0.01, *p < 0.05.

Our experiments yield a counterintuitive result (**Supplementary Table 5, 6**) with a minimal improvement in performance from using dense labels, yielding at best a 1%-2% statistically significant increase in median Dice score despite requiring an order of magnitude more labels. While previous experiments demonstrated that CryoViT benefited from training on more tomograms, additional annotations within the same tomograms failed to deliver comparable benefits, likely due to the high correlation between nearby slices in a tomogram. When coupled with nearly identical segmentation masks, the training data would be flooded with redundant input-output pairs. In contrast, including more tomograms in the training set would add new non-redundant information, explaining the improvements previously observed. Consequently, CryoViT trained on a select fraction of annotated slices matches the performance of an identical model trained on fully annotated tomograms, highlighting our method’s label efficiency.

## DISCUSSION

The development of generalizable deep learning models for cryoET data mining that can enable downstream quantitative analyses has been constrained by a lack of large-scale annotated datasets. Not only is it expensive to prepare and image samples using cryoET, but it is also difficult to manually annotate enough tomograms to build conventional U-Net-based segmentation models that accurately capture organelle morphology. Several strategies aimed at automating the annotation of subcellular features in cryoET tomograms have been proposed ^21^; however, most methods have relied on trained 2D U-Net models that consider a limited local context and therefore might not generalize well across tomograms. Furthermore, the emphasis of most prior methods has been overwhelmingly on annotation of membranes ^42,43^ or on localizing macromolecules ^44,45,46,47^ with the goal of deriving higher resolution structures through STA^3,48^. For instance, DeepFinder^44^ utilizes 3D U-Nets for localizing and segmenting macromolecules in cellular contexts, and DeepPiCt^46^ further improves upon this strategy by also localizing specific organelles including mitochondria and the endoplasmic reticulum, thus reducing false positives and restricting focus to biologically interesting regions. However, since these methods are designed specifically to detect small, relatively homogenous particles, their suitability for segmenting larger organelles is limited, especially in data- and label-constrained scenarios.

In response to these challenges, we developed CryoViT, a powerful method that enables accurate quantitative measurements of the morphology of pleomorphic organelles such as mitochondria. CryoViT leverages the implicit semantic information captured by vision foundation models to create accurate, data- and label-efficient segmentation systems that are robust across domain shifts induced by variations in phenotypes or other factors. Despite the fact that the ViT backbone powering CryoViT has not been trained on cryoET images but rather on over a hundred million natural images scraped from the internet ^49^, it nonetheless generates a rich set of features for tomogram slices that bear little resemblance to the original training data. This shows that by being exposed to a large and diverse array of images, the backbone possesses a surprisingly good general understanding of shapes, objects, and textures, which enables the construction of the efficient models we demonstrate here. Furthermore, our use of convolution kernels with a large depth dilation factor innovatively overcomes fundamental limitations of 2D vision foundation models. Our approach is demonstrated on a large, diverse dataset of 256 tomograms of mitochondria spanning 11 unique conditions representing a combination of healthy, diseased, and genetically modified samples derived from both human iPSCs and mouse models. Our dataset is the largest of its kind, resulting from hundreds of hours of annotation effort to generate “ground truth” labels, thereby enabling the training and evaluation of deep learning models.

Just as the introduction of CNNs for cryoET analysis ^21,50^ sparked the development of numerous methods for automated particle picking^44^, subtomogram averaging ^51,52,53^, and improved structural visualization by filling in the missing wedge^54^, including strategies leveraging alternative network models ^55^, vision foundation models hold great promise for large-scale quantitative analyses of pleomorphic structures in tomograms. As such, they represent a paradigm shift in how cryoET will be used to further our understanding of fundamental biological processes. Indeed, recent commentary ^56^ has highlighted the potential of foundational models for segmentation tasks, projecting that their accuracy and efficiency will make them the dominant tool for quantitative morphological studies. Other work ^57^ broadcasts a call-to-action to prioritize the development of automated methods to enable disease diagnosis and monitor disease progression by quantifying organelles, specifically mitochondria, in images from volume electron microscopy and related techniques that preserve specimens through conventional ^58,59^ or cryogenic ^6^ fixation. The precise segmentation masks generated by our approach can be used to quantitatively characterize phenotypic differences across healthy, disease-related, and drug-treated samples, making CryoViT an efficient and versatile tool that can yield fundamental biological and biomedical insights, including helping to validate the efficacy and safety of potential therapeutics.

As early work at the intersection of foundation models and cryoET, we envision that CryoViT will inspire further research in this rapidly evolving domain. To accelerate the development of the next generation of cryoET models, we provide an open-source software package for researchers to segment mitochondria with our pretrained models or build their own for other unique needs focusing on any structural targets detectable in cryoET datasets. Extensions of CryoViT will investigate modifications to our protocol, such as using self-supervised fine-tuning of the foundation model backbone to further improve feature extraction, as well as novel applications to study a diverse array of structures across various organisms, disease models, and patient samples, thereby advancing our understanding of cellular mechanisms and their implications in health and disease.

## ONLINE METHODS

### Tomogram preprocessing and annotation generation

Our analyses here were performed on previously published cryoET datasets of HD-patient-derived iPSCs differentiated into neurons, as well as HD mouse model neurons ^8^. Briefly, to clarify the properties of these tomograms, the motion-corrected tilt series were reconstructed with IMOD, smoothed with a SIRT-like filter set to 16 iterations to reduce noise and increase contrast, and low-pass filtered to 100 Å. To improve the contrast of features of interest further, the dynamic range for the tomograms’ intensity values was threshold filtered to ±3 standard deviations from the mean. This was followed by a linear rescaling to yield densities lying between -1.0 and 1.0 to standardize the input range for our CryoViT algorithm. The tomograms were further binned by a factor of 2 and clipped to produce uniform volumes 512 x 512 x 128 (height x width x depth) or 720 x 512 x 128, depending on whether the K2 or K3 camera was used to collect the corresponding tilt series. This step was necessary in order to train 3D U-Nets and CryoViT on entire tomograms without resorting to using patch-based approaches, given the constraints of limited GPU memory (NVIDIA A100-40GB).

From each tomogram, five slices containing mitochondria were selected from the binned-by-4 tomograms for annotation. The slices were selected so that each had at least one clearly visible mitochondrion, or a part of one, with a clearly visible double membrane separating it from the background. To span as much of the depth of the tomogram as possible, the extreme slices were chosen such that mitochondria were not clearly visible beyond them. The spacing between slices was kept uniform within each tomogram, but varied across tomograms based on how the mitochondria were distributed. The slices were uploaded to hasty.ai, a cloud annotation platform for manual labeling. After each annotation, a second annotator would independently verify the labels and correct any minor errors.

During the slice selection process, annotators also identified the z-limits, regions at either end of the tomogram beyond which they were certain that no mitochondria could be observed, either wholly or partially. The slices from the regions beyond the mitochondria were automatically labeled as fully background slices and were used as negative training references that complemented the five manually annotated slices at no additional annotation cost. Combined, these annotations formed the ground truth labels, with background voxels and foreground voxels (mitochondria) being marked with 0 and 1, respectively. The remaining unlabeled slices were marked with -1 so that predictions on these slices could be easily identified and ignored while training the model in a semi-supervised fashion.

### Feature extraction from vision transformers

Vision foundation models are designed to handle 2D RGB images; thus, features were independently extracted for each 2D slice along the depth dimension of tomograms. Since each 2D image is a single channel grayscale image, the values were repeated three times along a new channel axis to change the representation to an RGB image. Pixel intensity values (floating point numbers lying between 0 and 1) were normalized during the pre-training procedure of the underlying foundation model by subtracting the mean and dividing by the standard deviation of the RGB values, which are (0.485, 0.456, 0.406) and (0.229, 0.224, 0.225) for the DINOv2 family of foundation models, respectively.

Foundation models based on the vision transformer architecture generate embeddings for non-overlapping patches (usually 16 x 16 pixels in size) of an input RGB image ^29^. The output patch embeddings are then fed into the CryoViT model, which progressively upscales them by a factor of 2, repeated 4 times, to result in an output prediction of the same size as the original input image. However, DINOv2 foundation models use a slightly different patch size of 14 x 14 pixels, which is difficult to factorize into a sequence of hierarchical upscaling steps. Here, we implemented a simple workaround by scaling the height and width of the input image by a factor of ⅞, which yielded the same number of patches as using a patch size of 16 x 16 pixels on the original image.

The features extracted from each 2D slice were then stacked together along a new axis to create the overall features representation for each tomogram. As a concrete example, consider a tomogram with shape 512 x 512 x 128 (y, x, z). After preprocessing, each RGB slice of shape 3 x 512 x 512 (channels, x, y) is first be rescaled to 3 x 448 x 448 using bicubic interpolation, then fed into the DINOv2-g/14 foundation model, which yields a resulting “feature vector” of shape 1536 x 32 x 32 (channels x height x width) since the model produces patch embeddings with 1536 dimensions and there are 32 x 32 patches of 14 x 14 pixels in size in each 448 x 448 2D slice. Finally, the feature vectors for all slices (n=128 in this example) would be stacked together to create the feature vector of shape 1536 x 32 x 32 x 128 (channels x height x width x depth). Note that the depth dimension (z=128) remains unchanged and only the height and width (x,y) dimensions are downscaled by a factor of the patch size (14x, in this example).

### CryoViT model architecture

The CryoViT model is responsible for converting raw features extracted from the DINOv2 foundation model and decoding them into segmentation masks for the structures of interest (mitochondria in this case). It is composed of three components:

1. A fully connected layer to project the DINOv2 features from a 1536 dimensional space to a lower, 1024 dimensional space. This projection reduces the memory footprint of the tomogram’s features, enabling the analysis of full-size tomograms without breaking them up into smaller patches.
2. A sequence of four upscaling modules, each of which progressively upscales the input feature map by a factor of two in x and y. Collectively, they restore the feature map to its original resolution.
3. A non-linear output head which operates on the original-resolution feature map and produces voxel-wise predictions for the structure being segmented.

Each upscaling module, hereon referred to as a **synthesis block**, consists of two 3D convolution layers followed by a transposed 3D convolution layer. The convolution layers have anisotropic 3 x 3 x 3 convolution kernels, with the anisotropy arising from dilations in z (the “depth” dimension). Dilated convolutions ^37^ have been successfully used to model long-range feature interactions across multiple domains, such as very long DNA sequences ^60^. As part of the synthesis block, they primarily serve to aggregate information from feature maps spanning the depth of the tomogram. DINOv2 features are computed independently for each depthwise slice of the tomogram, so convolution kernels with a large depth dilation factor are vital for overcoming this fundamental limitation of 2D vision foundation models. The transposed convolution layer plays a complementary role and is responsible for upscaling the height and width (x,y) of the feature map of each slice by a factor of two. This is achieved with an anisotropic kernel of shape 2 x 2 x 1 (height x width x depth) with a stride of 2 x 2 x 1 since no upscaling is performed in the depth dimension.

The dilation rate and the number of convolution kernels in each layer in the synthesis block are carefully chosen to limit the memory footprint of the feature map as it is propagated through the CryoViT model. Since each upscaling stage results in an output feature map that is four times as large as its input, the number of channels in the feature map needs to be progressively reduced to avoid out-of-memory errors, even with high memory GPUs (NVIDIA A100-40GB). The CryoViT model aggressively decreases the number of channels in the feature map from 1024 to 8 over the span of the four synthesis blocks, inspired by the synthesis blocks used in U-Nets (see **Supplementary Table 7** for further details). The dilation rate is also progressively decayed from 32 to 1 so that earlier convolution layers aggregate coarse information across distant slices and later convolution layers model more fine-grained relationships across nearby slices of the tomogram. After upscaling through the cascade of synthesis blocks, the low dimensional feature map is passed through an output head with two 3 x 3 x 3 convolutions, followed by a sigmoid activation function to convert the output logits into per-voxel probability scores for the predicted segmentation mask. The network used Gaussian Error Linear Units ^61^ as activation functions after each layer, along with GroupNorm layers ^62^ at the start of each synthesis block to stabilize training with batch sizes as small as 1 tomogram.

CryoViT uses features extracted from tomograms using the DINOv2 ViT-g/14 foundation model which has 1.1 billion frozen parameters. The CryoViT model itself is relatively small and lightweight, with just 8.4 million trainable parameters.

### Modified 3D U-Net architecture

The 3D U-Net is designed based on the recipe provided by the nn-UNet software package. Presently, nn-UNet ^36^ does not support semi-supervised learning with sparsely labeled inputs, a fundamental requirement for label-efficient training. Our implementation closely matches the nn-UNet design and matches the parameter count of the CryoViT model to facilitate a fair comparison. Like the standard 3D U-Net, it has three downsampling (analysis) blocks and three upsampling (synthesis) blocks with skip connections between analysis and synthesis blocks at each level. It also uses transposed 3D convolutions for upscaling feature maps in the synthesis pathway, but the 2 x 2 x 2 max pooling layer is replaced with a 3D convolution layer with a 2 x 2 x 2 kernel and a matching stride, similar to the nn-UNet implementation. We also use their instance normalization strategy to stabilize training; that is, InstanceNorm layers ^63^ are inserted between each convolution and activation layer. Like the traditional 3D U-Net, the number of convolution kernels are increased at each level and we use heuristics inspired by nn-UNet to aggressively increase them while simultaneously staying within the limits of GPU memory as well as keeping the parameter count (8.5 million parameters) in the same ballpark as that of the CryoViT model. A full breakdown of the architecture is presented in **Supplementary Table 8.**

### Model training

All models were built and trained using the PyTorch Lightning framework. The compute and memory footprints of models were reduced by using 16-bit float precision and using PyTorch’s model compilation tools to generate optimized static graphs of the CryoViT model architecture. With this setup, training CryoViT models took anywhere from 10 minutes for a small sample of 15 tomograms up to 3 hours for the entire dataset of 256 tomograms. All models were trained using a single NVIDIA A100-40GB GPU.

We trained all our models with a batch size of one tomogram of shape 512 x 512 x 128. In some instances, tomograms had a shape of 720 x 512 x128, which exceeded the memory limits of the GPU while training, so we cropped those larger tomograms to slightly smaller dimensions of 512 x 512 x 128 voxels to maintain a uniform input size, spanning the entire tomogram in most cases. This random crop was applied directly to the raw tomogram in the case of the 3D U-Net, but while training CryoViT models, the crop was applied to the extracted DINOv2 features and 32 x 32 patches were cropped out from 45 x 32 patch regions, corresponding to a crop of 512 x 512 pixels from a larger region of 720 x 512 pixels. Note that we did not apply any cropping during inference as the memory requirements are lower and we could obtain predictions on the full-size tomograms (128 x 720 x 512 voxels).

To mitigate training instabilities caused by the small batch size, we used group normalization with up to 8 groups for CryoViT models. In our preliminary experiments, we observed no benefit to applying random augmentations with either the 3D U-Net or CryoViT, so we opted to forego them for computational efficiency. Similar to nn-UNet, we limited training to 50 epochs for all models and experiments where one epoch is defined as one complete iteration over the training tomograms. We carefully monitored training progress and verified that models were being trained to convergence within our training budget without overfitting to the data. Model weights were subject to L2 regularization with a weight decay coefficient of 1e-3. To further improve model generalization, we used Stochastic Weight Averaging ^64^ (SWA) for the last 20% of training. SWA can be thought of as an efficient model ensembling method by maintaining a moving average of the model’s weights during this last phase of training, causing it to converge to a better minima. We used the AdamW ^65^ optimizer with learning rates of 1e-4 and 3e-3 for CryoViT and the 3D U-Net, respectively.

Models were trained in a semi-supervised approach using sparsely labeled volumes. The input is propagated through the model to obtain a segmentation prediction for all voxels. The predictions for the unlabeled regions of the tomogram are discarded and the loss is only computed on a masked subset of voxels which have ground truth annotations. We trained the models using only the Dice loss ^38^ as we found the addition of the cross-entropy loss to have a negligible impact on training. While evaluating the model’s performance, we measured the mean Dice score of the model’s predictions binarized to 0-1 values using a threshold of 0.5.

### Cross validation and performance evaluation

The small size of some of our samples precluded model evaluation using a standard train / test split. We instead opted to use various forms of cross validation for our experiments to make the most of the limited data and created ten roughly equal splits for each of the 11 samples in our dataset. In our experiments on individual samples, we would train a new model on nine out of ten splits and evaluate it on the held-out split by recording the Dice scores for each tomogram in this split. This process was repeated ten times with each split serving as the test split exactly once, resulting in a Dice score being recorded for every tomogram in that sample. In experiments probing the data efficiency of CryoViT, we held out one sample and trained on a subset of tomograms from the remaining eleven samples. We made use of the predefined 10-fold splits to train on various fractions of data - for example, to train on 30% of the data, we trained on tomograms of the first 3 splits of each of the eleven sample conditions. Then, the trained model was evaluated on all tomograms from the held out (previously unseen) sample condition. In contrast, our experiments on generalization across domain shifts did not use any cross validation since we used distinct sample conditions for training and evaluation. Models were trained on all tomograms from either the healthy set (Q18, Q20, Wild Type) or the diseased set (Q53, Q66, Q109, BACHD, dN17 BACHD) and evaluated on the other set.

In all our experiments, we evaluated model predictions using the sparse ground truth labels consisting of five human annotated slices and several negative reference slices at the extremes of the z-axis which do not contain any mitochondria. This included our experiments involving models trained on densely labeled tomograms. Like the models trained on the sparse labels, these models’ predictions were evaluated on exactly the same sparse ground truth labels to facilitate a fair comparison.

To determine if the difference in segmentation performance of two models was statistically significant, we conducted paired significance tests in all our experiments. When comparing performance on individual samples (including domain generalization experiments), we used a non-parametric test, specifically, the Wilcoxon signed-rank test, due to the limited sample sizes with n=10 as the minimum value. Since experiments exploring data efficiency had a larger sample size of n=256, we instead used a t-test to compute p-values. All significance tests were paired and one-sided with the alternative hypothesis testing if one model is better than the other. When comparing CryoViT and 3D U-Nets, the alternative hypothesis is that CryoViT outperforms the 3D U-Net. When comparing the difference between CryoViT models trained on sparse and dense annotations, the alternative hypothesis is that models trained on dense annotations outperform those trained on sparse annotations. If the p-value calculated from the one-sided paired significance test is less than 0.05, the alternative hypothesis is accepted, otherwise we fail to reject the null hypothesis.

### Automated conversion of sparse labels to dense labels

While CryoViT demonstrates excellent generalization performance on unseen samples, across domain shifts, and also data constrained scenarios, it may occasionally fail to produce satisfactory results on particularly challenging specimens, such as those corresponding to more severe phenotypes or lower-quality reconstructions. In such instances, CryoViT’s predictions can be improved by fine-tuning it on these tomograms using human-generated sparse annotations for just five slices.

In our preliminary investigations, we observed that CryoViT models were able to produce near-perfect predictions on the unlabeled regions of the tomograms they were trained on. With models trained in a semi-supervised fashion, there is the possibility that they may overfit to the labeled slices and fail to generalize to the unlabeled regions. However this was not the case with CryoViT models which learned to smoothly interpolate in between the 5 labeled slices of each training tomogram, converting a handful of 2D annotations into a dense 3D segmentation mask that matches the contours of the 2D masks.

We opted to leverage this excellent generalization property to create AI-assisted dense annotations for each tomogram using the labeled regions as a guide for the model. To build a robust model less susceptible to overfitting, we trained a single CryoViT model on the entire dataset of 256 tomograms, and then applied it to make predictions on the same 256 tomograms. This trained model (also referred to as an “oracle”) is a proxy for human annotators and makes predictions for the same tomograms it was trained on. Next, we manually inspected each tomogram to verify if the dense segmentation prediction was consistent with the ground truth. In each case, the AI generated labels perfectly matched the ground truth, and in some cases, corrected minor errors in the manual ground truth annotations as well. We used these dense 3D predictions as training labels for our experiments on label efficiency. Note that the oracle model has no further role to play after generating the dense labels.

## DATA ACCESS AND CODE AVAILABILITY STATEMENT

The code and training models used in this study are available at https://github.com/sanketx/CryoVIT while the reconstructed tomograms and corresponding annotations will be made available through the EMPIAR database. We provide 256 tomograms spanning 11 sample conditions as compressed HDF5 files. These include sparse ground truth mitochondria annotations as well as dense annotations generated by CryoViT. The CryoViT software package can be used to reproduce our results or train and fine-tune models on custom datasets. We also provide trained models for mitochondria segmentation which can be used off-the-shelf.

## ACKNOWLEDGEMENTS

Special thank you to Dr. Charlene Smith-Geater (UC Irvine), Prof. Leslie Thompson (UC Irvine), Prof. William Mobley (UC San Diego), Prof. Chengbiao Wu (UC San Diego) and Prof. Judith Frydman (Stanford University) for their collaboration on past and ongoing projects related to our analyses.

## AUTHOR CONTRIBUTIONS

Conceptualization & methodology, SRG, SYL; software, SRG; annotation and validation, CH, GHW, JGGM; formal analysis, SRG; draft writing SRG, JGGM, WC; funding acquisition, SYL, WC. All authors have read the manuscript, provided feedback, and agreed to the published version.

