## Supplementary Information for "CryoViT: Efficient Segmentation of Cryogenic Electron Tomograms with Vision Foundation Models"

**Supplementary Table 1. Dataset composition by sample.**

| Sample | Number of Tomograms |
|---|---|
| Q53 PIAS1 | 10 |
| Q66 | 10 |
| Q53 | 14 |
| dN17 BACHD | 15 |
| Q66 GRFS1 | 17 |
| Q20 | 20 |
| Q18 | 21 |
| BACHD | 22 |
| Wild Type | 31 |
| Q109 | 33 |
| Q66 PIAS1 | 63 |
| **Total** | **256** |

**Supplementary Table 2. Model generalization on individual samples.**

| Sample | Median Dice Score | | Mean Dice Score ± Std | | Quartiles (Q1 - Q3) | | p-value |
|---|---|---|---|---|---|---|---|
| | CryoViT | 3D U-Net | CryoViT | 3D U-Net | CryoViT | 3D U-Net | |
| Q53 PIAS1 | 0.84 | 0.37 | 0.77 ± 0.22 | 0.43 ± 0.27 | 0.75 - 0.90 | 0.22 - 0.67 | 9.77E-04 |
| Q66 | 0.72 | 0.41 | 0.71 ± 0.18 | 0.49 ± 0.24 | 0.59 - 0.86 | 0.37 - 0.67 | 6.84E-03 |
| Q53 | 0.95 | 0.87 | 0.92 ± 0.06 | 0.87 ± 0.06 | 0.92 - 0.96 | 0.83 - 0.92 | 2.62E-03 |
| dN17 BACHD | 0.78 | 0.5 | 0.77 ± 0.15 | 0.45 ± 0.21 | 0.70 - 0.90 | 0.29 - 0.62 | 3.05E-05 |
| Q66 GRFS1 | 0.96 | 0.86 | 0.94 ± 0.05 | 0.82 ± 0.17 | 0.94 - 0.97 | 0.77 - 0.94 | 7.63E-06 |
| Q20 | 0.94 | 0.81 | 0.92 ± 0.06 | 0.75 ± 0.21 | 0.90 - 0.96 | 0.70 - 0.90 | 9.54E-07 |
| Q18 | 0.95 | 0.88 | 0.90 ± 0.14 | 0.82 ± 0.16 | 0.89 - 0.97 | 0.75 - 0.92 | 1.08E-03 |
| BACHD | 0.91 | 0.74 | 0.86 ± 0.13 | 0.72 ± 0.16 | 0.80 - 0.96 | 0.62 - 0.82 | 1.28E-04 |
| Wild Type | 0.95 | 0.88 | 0.92 ± 0.07 | 0.80 ± 0.20 | 0.88 - 0.97 | 0.80 - 0.93 | 1.11E-06 |
| Q109 | 0.97 | 0.87 | 0.93 ± 0.07 | 0.80 ± 0.21 | 0.92 - 0.98 | 0.77 - 0.94 | 3.57E-08 |
| Q66 PIAS1 | 0.96 | 0.93 | 0.91 ± 0.13 | 0.87 ± 0.15 | 0.92 - 0.97 | 0.87 - 0.95 | 1.58E-05 |

**Supplementary Table 3. Model generalization across domain shifts.**

| Sample | Median Dice Score | | Mean Dice Score ± Std | | Quartiles (Q1 - Q3) | | p-value |
|---|---|---|---|---|---|---|---|
| | CryoViT | 3D U-Net | CryoViT | 3D U-Net | CryoViT | 3D U-Net | |
| Q20 | 0.94 | 0.85 | 0.91 ± 0.05 | 0.85 ± 0.10 | 0.87 - 0.96 | 0.79 - 0.93 | 1.61E-04 |
| Q18 | 0.96 | 0.85 | 0.92 ± 0.13 | 0.83 ± 0.15 | 0.92 - 0.97 | 0.81 - 0.92 | 4.26E-04 |
| Wild Type | 0.95 | 0.77 | 0.91 ± 0.13 | 0.74 ± 0.21 | 0.90 - 0.98 | 0.62 - 0.93 | 6.38E-08 |
| Q66 | 0.66 | 0.41 | 0.65 ± 0.28 | 0.41 ± 0.28 | 0.47 - 0.90 | 0.17 - 0.57 | 9.77E-03 |
| Q53 | 0.94 | 0.85 | 0.92 ± 0.07 | 0.83 ± 0.09 | 0.90 - 0.97 | 0.79 - 0.88 | 6.10E-05 |
| dN17 BACHD | 0.82 | 0.39 | 0.75 ± 0.20 | 0.38 ± 0.24 | 0.65 - 0.89 | 0.21 - 0.54 | 9.16E-05 |
| BACHD | 0.95 | 0.79 | 0.80 ± 0.28 | 0.76 ± 0.20 | 0.84 - 0.97 | 0.69 - 0.89 | 6.03E-02 |
| Q109 | 0.9 | 0.91 | 0.90 ± 0.06 | 0.81 ± 0.24 | 0.85 - 0.95 | 0.80 - 0.95 | 1.54E-02 |

**Supplementary Table 4. Data efficiency and model generalization on unseen samples.**

| % of Training Data | Median Dice Score | | Mean Dice Score ± Std | | Quartiles (Q1 - Q3) | | p-value |
|---|---|---|---|---|---|---|---|
| | CryoViT | 3D U-Net | CryoViT | 3D U-Net | CryoViT | 3D U-Net | |
| 10% | 0.91 | 0.75 | 0.85 ± 0.16 | 0.64 ± 0.27 | 0.82 - 0.95 | 0.46 - 0.85 | 1.16E-36 |
| 20% | 0.93 | 0.8 | 0.87 ± 0.15 | 0.70 ± 0.26 | 0.87 - 0.96 | 0.58 - 0.89 | 3.05E-31 |
| 30% | 0.95 | 0.82 | 0.89 ± 0.14 | 0.71 ± 0.26 | 0.88 - 0.97 | 0.62 - 0.90 | 2.64E-29 |
| 40% | 0.95 | 0.84 | 0.90 ± 0.13 | 0.73 ± 0.25 | 0.89 - 0.97 | 0.60 - 0.91 | 1.19E-28 |
| 50% | 0.95 | 0.83 | 0.90 ± 0.12 | 0.74 ± 0.24 | 0.89 - 0.97 | 0.66 - 0.92 | 1.08E-28 |
| 60% | 0.95 | 0.84 | 0.91 ± 0.12 | 0.76 ± 0.23 | 0.90 - 0.97 | 0.64 - 0.93 | 7.43E-27 |
| 70% | 0.95 | 0.86 | 0.91 ± 0.11 | 0.77 ± 0.23 | 0.91 - 0.97 | 0.68 - 0.93 | 3.61E-24 |
| 80% | 0.96 | 0.88 | 0.91 ± 0.12 | 0.79 ± 0.22 | 0.91 - 0.97 | 0.73 - 0.94 | 1.89E-20 |
| 90% | 0.95 | 0.87 | 0.91 ± 0.12 | 0.79 ± 0.22 | 0.90 - 0.97 | 0.72 - 0.95 | 1.14E-19 |
| 100% | 0.96 | 0.89 | 0.91 ± 0.12 | 0.82 ± 0.20 | 0.90 - 0.97 | 0.77 - 0.95 | 2.89E-16 |

**Supplementary Table 5. Impact of sparse vs dense training labels on individual samples.**

| Sample | Median Dice Score | | Mean Dice Score ± Std | | Quartiles (Q1 - Q3) | | p-value |
|---|---|---|---|---|---|---|---|
| | Sparse | Dense | Sparse | Dense | Sparse | Dense | |
| Q53 PIAS1 | 0.84 | 0.87 | 0.77 ± 0.22 | 0.78 ± 0.23 | 0.75 - 0.90 | 0.76 - 0.92 | 1.61E-01 |
| Q66 | 0.72 | 0.79 | 0.71 ± 0.18 | 0.74 ± 0.17 | 0.59 - 0.86 | 0.63 - 0.85 | 1.16E-01 |
| Q53 | 0.95 | 0.96 | 0.92 ± 0.06 | 0.94 ± 0.05 | 0.92 - 0.96 | 0.93 - 0.97 | 1.83E-04 |
| dN17 BACHD | 0.78 | 0.79 | 0.77 ± 0.15 | 0.77 ± 0.15 | 0.70 - 0.90 | 0.67 - 0.92 | 2.81E-01 |
| Q66 GRFS1 | 0.96 | 0.95 | 0.94 ± 0.05 | 0.95 ± 0.04 | 0.94 - 0.97 | 0.94 - 0.97 | 3.06E-01 |
| Q20 | 0.94 | 0.93 | 0.92 ± 0.06 | 0.93 ± 0.05 | 0.90 - 0.96 | 0.90 - 0.97 | 1.83E-03 |
| Q18 | 0.95 | 0.95 | 0.90 ± 0.14 | 0.91 ± 0.10 | 0.89 - 0.97 | 0.88 - 0.97 | 1.87E-01 |
| BACHD | 0.91 | 0.92 | 0.86 ± 0.13 | 0.88 ± 0.11 | 0.80 - 0.96 | 0.83 - 0.97 | 2.34E-03 |
| Wild Type | 0.95 | 0.96 | 0.92 ± 0.07 | 0.92 ± 0.08 | 0.88 - 0.97 | 0.87 - 0.98 | 1.94E-01 |
| Q109 | 0.97 | 0.97 | 0.93 ± 0.07 | 0.94 ± 0.07 | 0.92 - 0.98 | 0.94 - 0.98 | 1.36E-03 |
| Q66 PIAS1 | 0.96 | 0.96 | 0.91 ± 0.13 | 0.92 ± 0.12 | 0.92 - 0.97 | 0.93 - 0.97 | 1.08E-06 |

**Supplementary Table 6. Impact of sparse vs dense training labels on data efficiency and model generalization on unseen samples.**

| % of Training Data | Median Dice Score | | Mean Dice Score ± Std | | Quartiles (Q1 - Q3) | | p-value |
|---|---|---|---|---|---|---|---|
| | Sparse | Dense | Sparse | Dense | Sparse | Dense | |
| 10% | 0.91 | 0.92 | 0.85 ± 0.16 | 0.86 ± 0.15 | 0.82 - 0.95 | 0.83 - 0.96 | 3.55E-03 |
| 20% | 0.93 | 0.94 | 0.87 ± 0.15 | 0.89 ± 0.14 | 0.87 - 0.96 | 0.89 - 0.97 | 5.87E-08 |
| 30% | 0.95 | 0.95 | 0.89 ± 0.14 | 0.90 ± 0.13 | 0.88 - 0.97 | 0.90 - 0.97 | 7.46E-05 |
| 40% | 0.95 | 0.96 | 0.90 ± 0.13 | 0.91 ± 0.12 | 0.89 - 0.97 | 0.91 - 0.97 | 1.57E-03 |
| 50% | 0.95 | 0.96 | 0.90 ± 0.12 | 0.91 ± 0.12 | 0.89 - 0.97 | 0.91 - 0.97 | 1.01E-02 |
| 60% | 0.95 | 0.96 | 0.91 ± 0.12 | 0.92 ± 0.11 | 0.90 - 0.97 | 0.92 - 0.97 | 9.48E-05 |
| 70% | 0.95 | 0.96 | 0.91 ± 0.11 | 0.92 ± 0.10 | 0.91 - 0.97 | 0.92 - 0.98 | 4.16E-04 |
| 80% | 0.96 | 0.96 | 0.91 ± 0.12 | 0.92 ± 0.11 | 0.91 - 0.97 | 0.92 - 0.98 | 2.76E-04 |
| 90% | 0.95 | 0.96 | 0.91 ± 0.12 | 0.93 ± 0.10 | 0.90 - 0.97 | 0.92 - 0.98 | 2.22E-08 |
| 100% | 0.96 | 0.96 | 0.91 ± 0.12 | 0.93 ± 0.10 | 0.90 - 0.97 | 0.93 - 0.98 | 2.55E-07 |

**Supplementary Table 7. CryoViT Model Architecture.**
All convolution and projection layers are followed by a GELU activation layer. Each synthesis block is preceded by a group normalization layer with 8 groups.

| Layer | Output Shape | Notes |
|---|---|---|
| Resize (preprocessing) | (1, 128, 448, 448) (C, D, H, W) | Height and width are resized from 512 x 512. |
| DINOv2 features (precomputed) | (1536, 128, 32, 32) (C, D, H, W) | 14 x 14 patches are vectorized in the H, W dimensions. The height and width are downscaled from 448 x 448 -> 32 x 32. |
| Linear Projection | (1024, 128, 32, 32) (C, D, H, W) | Linear projection to fewer dimensions to reduce memory footprint. Channels: 1536 -> 1024 |
| **Synthesis Block 0** | | |
| 3D Convolution | (192, 128, 64, 64) (C, D, H, W) | Dilated 3D convolution with a 3 x 3 x 3 anisotropic kernel. Channels: 1024 -> 192. Depth dilation: 32 |
| 3D Convolution | (192, 128, 64, 64) (C, D, H, W) | Dilated 3D convolution with a 3 x 3 x 3 anisotropic kernel. Channels: 192 -> 192. Depth dilation: 24 |
| 2D Upsampling (upscale by 2) | (128, 128, 64, 64) (C, D, H, W) | Transposed 3D convolution with a kernel and stride of 1 x 2 x 2 Channels: 192 -> 128 |
| **Synthesis Block 1** | | |
| 3D Convolution | (64, 128, 64, 64) (C, D, H, W) | Dilated 3D convolution with a 3 x 3 x 3 anisotropic kernel. Channels: 128 -> 64. Depth dilation: 16 |
| 3D Convolution | (64, 128, 64, 64) (C, D, H, W) | Dilated 3D convolution with a 3 x 3 x 3 anisotropic kernel. Channels: 64 -> 64. Depth dilation: 12 |
| 2D Upsampling (upscale by 2) | (32, 128, 128, 128) (C, D, H, W) | Transposed 3D convolution with a kernel and stride of 1 x 2 x 2 Channels: 64 -> 32 |
| **Synthesis Block 2** | | |
| 3D Convolution | (32, 128, 128, 128) (C, D, H, W) | Dilated 3D convolution with a 3 x 3 x 3 anisotropic kernel. Channels: 32 -> 32. Depth dilation: 8 |
| 3D Convolution | (32, 128, 128, 128) (C, D, H, W) | Dilated 3D convolution with a 3 x 3 x 3 anisotropic kernel. Channels: 32 -> 32. Depth dilation: 4 |
| 2D Upsampling (upscale by 2) | (32, 128, 256, 256) (C, D, H, W) | Transposed 3D convolution with a kernel and stride of 1 x 2 x 2 Channels: 32 -> 32 |
| **Synthesis Block 3** | | |
| 3D Convolution | (16, 128, 256, 256) (C, D, H, W) | Dilated 3D convolution with a 3 x 3 x 3 anisotropic kernel. Channels: 32 -> 16. Depth dilation: 2 |
| 3D Convolution | (16, 128, 256, 256) (C, D, H, W) | Regular 3D convolution with a 3 x 3 x 3 anisotropic kernel. Channels: 16 -> 16. |

| | | |
|---|---|---|
| 2D Upsampling (upscale by 2) | (8, 128, 512, 512) (C, D, H, W) | Transposed 3D convolution with a kernel and stride of 1 x 2 x 2<br>Channels: 16 -> 8 |
| **Output Layer** | | |
| 3D Convolution | (8, 128, 512, 512) (C, D, H, W) | 3 x 3 x 3 kernel, zero padded to preserve output size.<br>Channels: 8 -> 8 |
| 3D Convolution | (1, 128, 512, 512) (C, D, H, W) | Predicted logits for each voxel with a 3 x 3 x 3 kernel.<br>Channels: 8 -> 1 |

**Supplementary Table 8. Modified 3D U-Net Model Architecture.**
All convolution and projection layers are followed by an instance normalization layer and a GELU activation layer.

| Layer | Output Shape | Notes |
|---|---|---|
| Pad Input (preprocessing) | (1, 128, 512, 512) (C, D, H, W) | Height, width, and depth are zero padded to be multiples of 16.<br>Required for inputs of arbitrary size. |
| **Analysis Block 0** | | |
| 3D Convolution | (16, 128, 512, 512) (C, D, H, W) | 3 x 3 x 3 kernel, zero padded to preserve output size.<br>Channels: 1 -> 16 |
| 3D Convolution (skip connection) | (16, 128, 512, 512) (C, D, H, W) | 3 x 3 x 3 kernel, zero padded to preserve output size.<br>Channels: 16 -> 16 |
| 3D Pooling (downscale by 2) | (16, 64, 256, 256) (C, D, H, W) | 3D convolution with a kernel and stride of 2 x 2 x 2.<br>Channels: 16 -> 16 |
| **Analysis Block 1** | | |
| 3D Convolution | (64, 64, 256, 256) (C, D, H, W) | 3 x 3 x 3 kernel, zero padded to preserve output size.<br>Channels: 16 -> 64 |
| 3D Convolution (skip connection) | (64, 64, 256, 256) (C, D, H, W) | 3 x 3 x 3 kernel, zero padded to preserve output size.<br>Channels: 64 -> 64 |
| 3D Pooling (downscale by 2) | (64, 32, 128, 128) (C, D, H, W) | 3D convolution with a kernel and stride of 2 x 2 x 2<br>Channels: 64 -> 64 |
| **Analysis Block 2** | | |
| 3D Convolution | (256, 32, 128, 128) (C, D, H, W) | 3 x 3 x 3 kernel, zero padded to preserve output size.<br>Channels: 64 -> 256 |
| 3D Convolution (skip connection) | (256, 32, 128, 128) (C, D, H, W) | 3 x 3 x 3 kernel, zero padded to preserve output size.<br>Channels: 256 -> 256 |
| 3D Pooling (downscale by 2) | (256, 16, 64, 64) (C, D, H, W) | 3D convolution with a kernel and stride of 2 x 2 x 2<br>Channels: 256 -> 256 |

| | | |
|---|---|---|
| **Bottom Block** | | |
| 3D Convolution | (384, 16, 64, 64)<br>(C, D, H, W) | 3 x 3 x 3 kernel, zero padded to preserve output size.<br>Channels: 256 -> 384 |
| 3D Convolution | (256, 16, 64, 64)<br>(C, D, H, W) | 3 x 3 x 3 kernel, zero padded to preserve output size.<br>Channels: 384 -> 256 |
| **Synthesis Block 0** | | |
| 3D Upsampling<br>(upscale by 2) | (64, 32, 128, 128)<br>(C, D, H, W) | Transposed 3D convolution with a kernel and stride of 2 x 2 x 2<br>Channels: 256 -> 64 |
| Concatenate | (320, 32, 128, 128)<br>(C, D, H, W) | Channelwise concatenation of the upsampled tensor and the skip connection from Analysis Block 2. |
| Linear Projection | (64, 32, 128, 128)<br>(C, D, H, W) | Linear projection to fewer dimensions to reduce memory footprint.<br>Channels: 320 -> 64 |
| 3D Convolution | (64, 32, 128, 128)<br>(C, D, H, W) | 3 x 3 x 3 kernel, zero padded to preserve output size.<br>Channels: 64 -> 64 |
| **Synthesis Block 1** | | |
| 3D Upsampling<br>(upscale by 2) | (16, 64, 256, 256)<br>(C, D, H, W) | Transposed 3D convolution with a kernel and stride of 2 x 2 x 2<br>Channels: 64 -> 16 |
| Concatenate | (80, 64, 256, 256)<br>(C, D, H, W) | Channelwise concatenation of the upsampled tensor and the skip connection from Analysis Block 1. |
| Linear Projection | (16, 64, 256, 256)<br>(C, D, H, W) | Linear projection to fewer dimensions to reduce memory footprint.<br>Channels: 80 -> 16 |
| 3D Convolution | (16, 64, 256, 256)<br>(C, D, H, W) | 3 x 3 x 3 kernel, zero padded to preserve output size.<br>Channels: 16 -> 16 |
| **Synthesis Block 2** | | |
| 3D Upsampling<br>(upscale by 2) | (16, 128, 512, 512)<br>(C, D, H, W) | Transposed 3D convolution with a kernel and stride of 2 x 2 x 2<br>Channels: 16 -> 16 |
| Concatenate | (32, 128, 512, 512)<br>(C, D, H, W) | Channelwise concatenation of the upsampled tensor and the skip connection from Analysis Block 0. |
| Linear Projection | (16, 128, 512, 512)<br>(C, D, H, W) | Linear projection to fewer dimensions to reduce memory footprint.<br>Channels: 32 -> 16 |
| 3D Convolution | (16, 128, 512, 512)<br>(C, D, H, W) | 3 x 3 x 3 kernel, zero padded to preserve output size.<br>Channels: 16 -> 16 |
| **Output Layer** | | |
| Linear Projection | (1, 128, 512, 512)<br>(C, D, H, W) | Predicted logits for each voxel.<br>Channels: 16 -> 1 |
